# Repeat-driven generation of antigenic diversity in a major human pathogen, *Trypanosoma cruzi*

**DOI:** 10.1101/283531

**Authors:** Carlos Talavera-López, João Luís Reis-Cunha, Louisa A. Messenger, Michael D. Lewis, Matthew Yeo, Daniella C. Bartholomeu, José E. Calzada, Azael Saldaña, Juan David Ramírez, Felipe Guhl, Sofía Ocaña-Mayorga, Jaime A. Costales, Rodion Gorchakov, Kathryn Jones, Melissa Nolan Garcia, Edmundo C. Grisard, Santuza M. R. Teixeira, Hernán Carrasco, Maria Elena Bottazzi, Peter J. Hotez, Kristy O. Murray, Mario J. Grijalva, Barbara Burleigh, Michael A. Miles, Björn Andersson

## Abstract

*Trypanosoma cruzi*, a zoonotic kinetoplastid protozoan parasite with a complex genome, is the causative agent of American trypanosomiasis (Chagas disease). Having a very plastic and complex genome, the parasite uses a highly diverse repertoire of surface molecules, with pivotal roles in cell invasion, immune evasion and pathogenesis. Thus far, the complexity of the genomic regions containing these genes has impaired the assembling a genome at chromosomal level, turning impossible to study the structure and function of the several thousand repetitive genes encoding the surface molecules of the parasite. We here present the genome assembly of a *T. cruzi* clade I (TcI) strain at semi-chromosomal level using high coverage PacBio single molecule sequencing, together with whole genome Illumina sequencing of 34 *T. cruzi* TcI isolates and clones from different geographic locations, sample sources and clinical outcomes. Resolution of the surface molecule gene distribution reveals an unusual duality in the organisation of the parasite genome, a synteny of the core genomic region with related protozoa flanked by unique and highly plastic multigene families’ clusters encoding surface antigens. The presence of abundant interspersed retrotransposons in these multigene families’ clusters suggests that these elements are involved in a recombination mechanism for the generation of antigenic variation and evasion of the host immune response on these TcI strains. The comparative genomic analysis of the cohort of TcI strains revealed multiple cases of such recombination events involving surface molecule genes and has provided new insights into *T. cruzi* population structure.

## INTRODUCTION

*Trypanosoma cruzi* is a kinetoplastid protozoan and the etiologic agent of Chagas disease, considered one of the most important human parasitic disease in Latin America. The Global Burden of Disease Study 2013 reported that almost 7 million people live with Chagas disease in the Western Hemisphere^1^, with the expectation that up to one third will progress to develop chronic chagasic cardiomyopathy (CCC) or other life-threatening symptoms. In 2015, 5,742,167 people were estimated to be infected with *T. cruzi* in 21 Latin American countries and around 13 % of the Latin American population is at risk of contracting *T. cruzi* infection due to domicile infestation of triatomine bugs or due to non-vectorial transmission via blood transfusion, organ transplant, oral, congenital or accidental infection^2^. Human Chagas disease is not restricted to Latin America. Migration of infected humans to non-endemic areas has made it a new public health threat in other geographic areas such as North America, Europe and Asia^4^. Also, sylvatic *T. cruzi* transmission cycles, often associated with human disease, have been described in areas formerly considered as free from this disease such as in Texas (USA)^4^.

The acute phase of the disease frequently lacks specific symptoms, is often undiagnosed and usually resolves in a few weeks in immunocompetent individuals but may be fatal in around 5% of diagnosed cases. Without successful treatment, a *T. cruzi* infection is normally carried for life. The disease progresses to either a chronic indeterminate phase that is asymptomatic, or to a chronic symptomatic phase with severe clinical syndromes such as cardiomyopathy, megaesophagus and/or megacolon^5^; meningoencephalitis may occur, especially in immunocompromised patients^4^. The current prolonged chemotherapy (benznidazole or nifurtimox) is mostly effective only in the acute phase, particularly because severe side effects may interrupt treatment of adults in the chronic phase. There is currently no effective treatment for advanced chagasic cardiomyopathy^6^, and there is an urgent need to identify new potential drug and vaccine targets^7^.

*T. cruzi* infection is a zoonosis, and the parasite has a complex life cycle; where transmission to humans occurs most frequently by contamination with infected feces from triatomine insect vectors (Subfamily Triatominae). The parasite evades the immune responses with the aid of multiple surface molecules from three large diverse gene families (Trans-Sialidases, Mucins and Mucin-Associated Surface Proteins - MASPs), which are also involved in cell invasion and possibly pathogenicity^8^.

Six distinct genetic clades of *T. cruzi* have been recognised, named TcI to TcVI (Discrete Typing Units or DTU-I to VI). The first genome sequence for *T. cruzi* was produced using Sanger sequencing technology from a hybrid, highly polymorphic, TcVI strain. The resultant genome sequence, while extremely useful for the core regions of the genome, was highly fragmented, especially in repetitive regions^9^. This sequence has been improved using enhanced scaffolding algorithms, but the repetitive regions remained unresolved^10^. Subsequently, FLX 454 Titanium and Illumina sequencing were used to sequence a less polymorphic TcI strain (Sylvio X10/1), which allowed the first comparative genomic studies of *T. cruzi*, but correct assembly of repetitive regions was still impossible^11,12^. The thousands of related genes that code for the surface proteins are generally located in large multigene families clusters of the *T. cruzi* genome^13^, in the form of extremely repetitive segments with multiple gene copies and pseudogenes. These multigene families’ clusters are distinct from the core regions of the genome in synteny, gene content and diversity^22^. The repetitive nature of the tandem arrays and the length of the repeat regions, have made correct assembly impossible using short and medium-sized sequence reads. The available *T. cruzi* genome sequences are therefore incomplete and erroneous in these important regions, making it impossible to study the complex surface gene families.

The population structure of *T. cruzi* is complex, and there is a high degree of genetic and phenotypic variation. The current TcI to TcVI clades are based on biochemical and molecular markers^14^, although there is substantial diversity even within these six groups^15^. The TcI clade is widespread and can be found across the American continent, and has been associated with CCC^16^ and sudden death^17,18^, among other clinical manifestations.

We have produced a complete and reliable reference sequence of the majority (∼98.5%) of the *T. cruzi* TcI Sylvio X10/1 genome, and we have generated Illumina whole genome sequencing data for 34 *T. cruzi* TcI isolates and clones from different geographic locations for comparative analyses. Thus, we have been able to decipher the majority of the organisation of *T. cruzi* surface protein coding gene repertoire from TcI Sylvio X10/1 strain, revealing large numbers of evenly spaced retrotransposons, which may play a role in generating genomic structural diversity and antigenic variation. Furthermore, the comparative data enabled the first exploration of whole-genome population genetics of *T. cruzi* in different environments and geographic locations. We found patterns of active recombination associated with generation of new surface molecule variants. Together, these results contribute to answering longstanding questions on the biology of Chagas disease and parasitism in general. The availability of the complete repertoire of genes encoding surface molecules allows further research on virulence and pathogenesis, as well as the identification of drug targets and vaccine candidates, focused on shared and conserved motifs present within these variable families.

## RESULTS

### Genome sequence of *Trypanosoma cruzi* Sylvio X10/1

The final Sylvio X10/1 (TcI) genome assembly reconstructed 98.5 % of the estimated strain genome size and was contained in 47 scaffolds (**Figure 1a**) - which here will be referred to as pseudomolecules - assembled from 210 X PacBio sequence data and a previous Illumina data set (**Table 1**). Comparison with the available assembly of the TcVI strain CL Brener revealed a conserved core of syntenic blocks composed of stretches of homologous sequences separated by large gaps of sequence that were not reconstructed in the hybrid TcVI strain. These gaps mostly corresponded to surface molecule gene arrays and simple repeats in the Sylvio X10/1 genome (**Figure 1b**). The length of the PacBio reads and the high coverage allowed the reconstruction of long stretches of repetitive sequences (**Figure 1b**) that could not be resolved using shorter read data.

**Figure 1:**
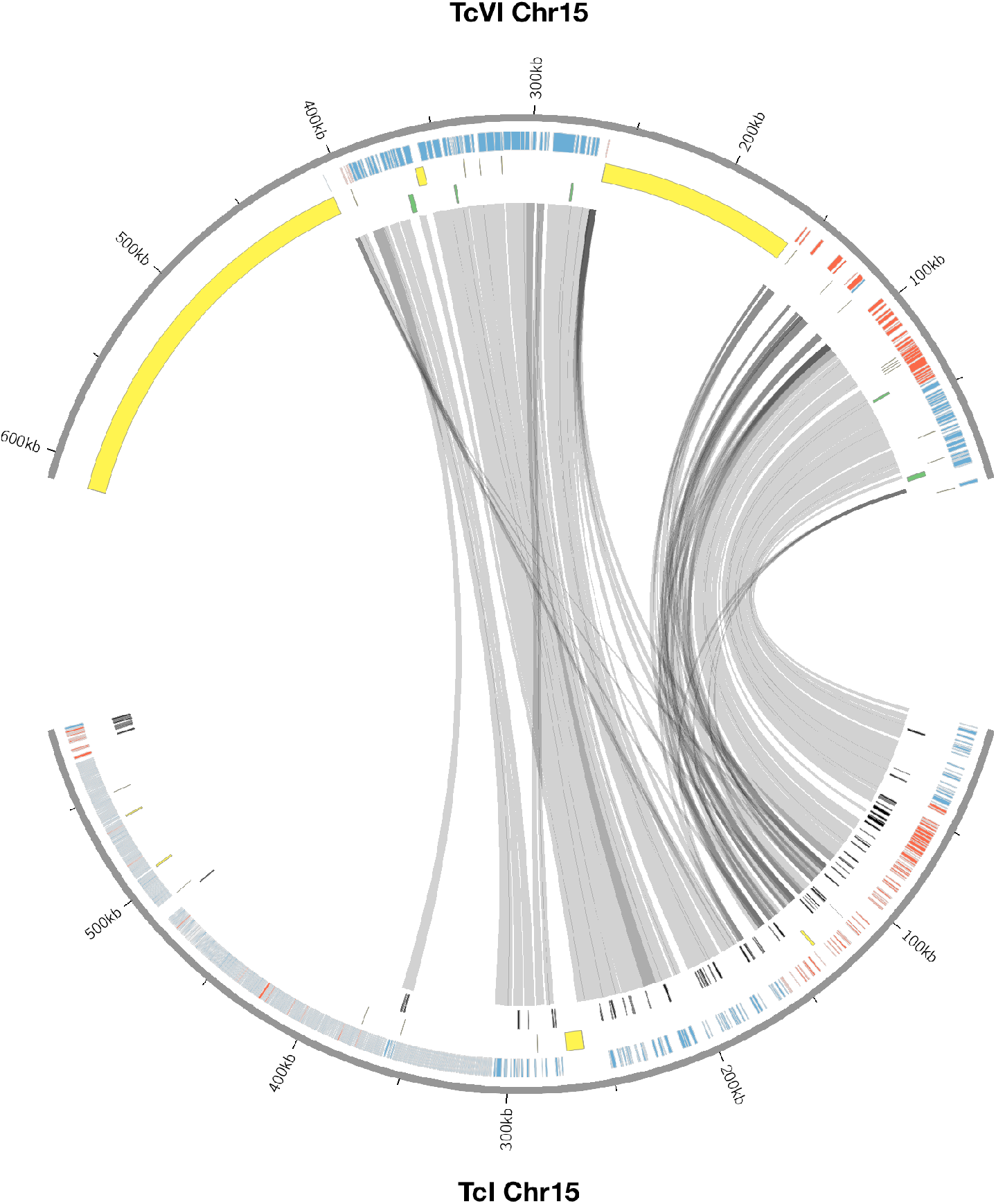
**a**) Distribution of surface molecule gene tandem arrays in the 47 chromosomes of the TcI Sylvio X10/1 genome. In this image, each line correspond to a pseudomolecule drawn in proportion to its size, where genes corresponding to the largest *T. cruzi* multigene families are represented by colored boxes, hypothetical genes are represented by grey boxes and the other genes as black boxes. The position of the gene box above or below correspond respectively to its presence in the + or − strand. **b**) Comparison of chromosome 15 from the TcVI CL Brener and TcI Sylvio X10/1 assemblies. Grey lines between pseudomolecules represent regions of synteny between orthologous genes, and yellow blocks represent gaps in both genome assemblies. Coloured blocks represent paralogs of multicopy gene families. **c**) Distribution of surface molecule gene tandem arrays in the 47 chromosomes of the TcI Sylvio X10/1 genome.

**Figure 2:**
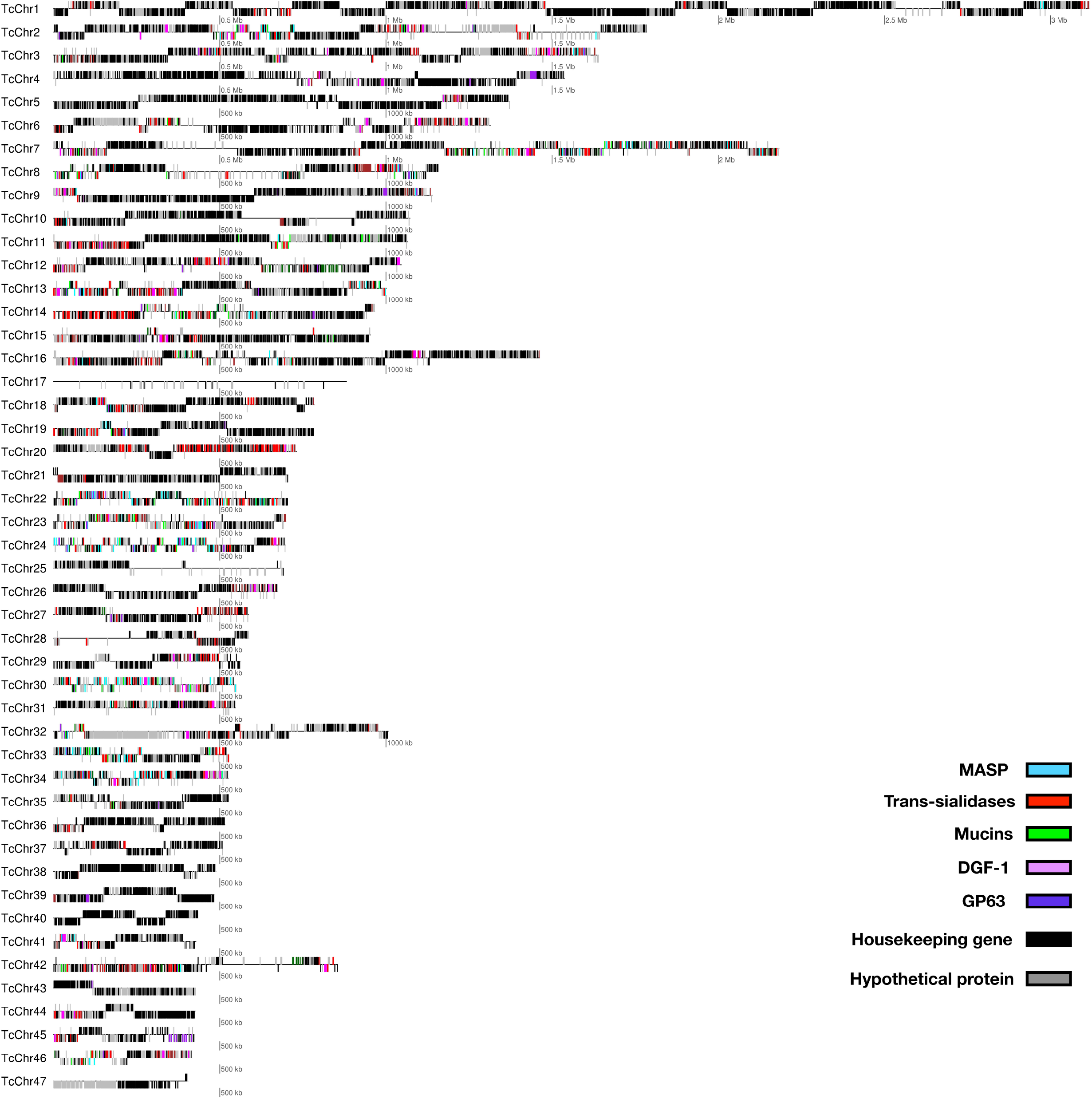
**a**) Bayesian principal component analysis (PCA) of *T. cruzi* TcI strains using INDELs and **b**) Identity by Descent (IBD) dendogram of *T. cruzi* TcI strains using SNPs. Both analyses, using different markers, support the population structure of the analysed TcI samples. Notably, the highly virulent TcI Colombiana and the Panamanian TcI H1 from a chronic patient are presented as outliers (**b**, far left).

**Table 1:** *Trypanosoma cruzi* Sylvio X10/1 strain (Tc-I) genome assembly.

| Metric | Value |
| --- | --- |
| Genome size | 41.3 Mbp |
| Number of scaffolds | 47 |
| Percentage of reconstruction | 98.5 % |
| Longest scaffold | 3.1 Mbp |
| Shortest scaffold | 404 Kb |
| NG50 | 1.0 Mbp |
| N50 | 1.1 Mbp |

The coverage of genomic regions coding for surface molecules supported the correct reconstruction of these areas (**Figure 1a**). To further investigate the quality of the new assembly, Illumina short reads were mapped and analysed with FRC_bam, which revealed assembly artefacts related to low coverage, wrong paired end read orientation, and higher than expected sequencing coverage in regions with long stretches of simple repeats.

Repetitive elements comprised 18.43 % of the TcI Sylvio X10/1 genome, 2.18 % of which cannot be classified using the repeat databases. LINE retroelements of the R1/Jockey group (3.63 %) and VIPER LTRs (2.87 %) were found to be the most prevalent types of retroelements, covering 6.89 % of the genome, which is much higher than the 2.57 % estimated from the previously published Sylvio X10/1 draft assembly using short reads^12^.

Although retrotransposons were found to be present throughout the genome, the frequency of VIPER and L1Tc elements was markedly higher in multigene families clusters regions and they were found within one kilobase of pseudogenes, hypothetical proteins and surface molecule gene tandem arrays (One-sided Fisher exact test, *p-value* < 1.32 e-16). This distribution indicates that these elements may play a role in the generation of new sequence diversity in gene families clusters by providing a source of microhomology. It is compelling to speculate that they could act as transcriptional regulators by the introduction of novel transcription start sites, as has been proposed in other eukaryotes^19^; nevertheless, we do not have experimental evidence for the activity of these retroelements in *T. cruzi*.

Simple and low complexity repeats were observed surrounding subtelomeric coding sequences and were also more abundant in the subtelomeric regions (2.18 %), extending up to 4 Kb, compared to core regions (0.98 %) where they were much shorter (10 - 120 bp). The most prevalent type of simple repeat had the (C)n motif (11.7 %), (TG)n repeat motif (5.6 %) and (CA)n repeat motif (5.1 %); each variable in length. This microhomology of the simple subtelomeric repeats may facilitate recombination for the generation of new surface molecule variants, as described in other parasitic protozoa, including *Trypanosoma brucei* and *Plasmodium falciparum*^20,21^. However, it is noteworthy to mention that such subtelomeric regions are far less complex and shorter in *T. brucei* (african, virulent) and *T. rangeli*^23^ (american, non-virulent).

Based on our annotation approach, a total of 19,096 genes were identified in the TcI Sylvio X10/1 haploid genome sequence, compared to the higher estimate of 22,570 for TcVI CL Brener, mostly due to the larger size of the multigene families’ clusters in the TcVI hybrid genome. The core regions of the genome were found to correspond well to results generated previously using short read sequencing of the same strain^11^, in both gene organisation and content. Tandem repeated genes that were collapsed in previous *T. cruzi* genome assemblies were now resolved. About 24.1% (n = 4,602) of the total annotated genes were truncated, mostly due to the introduction of premature stop codons, and 67 % of these were located within surface molecule gene arrays, sharing motifs of the complete genes.

The new assembly allowed an improved analysis of the *T. cruzi* surface molecule gene repertoire. Genes of each of the three major surface molecules families were organised as multiple tandem arrays. After genome annotation, the total number of such arrays were: trans-sialidases, 312, with 2,048 complete gene copies and 201 pseudogenes; mucins 98, with 2,466 complete copies and 111 pseudogenes; MASPs 264, with 1,888 complete copies and 245 pseudogenes. These three surface molecule gene families comprised 16.02 Mbp (39.04 %) of the TcI Sylvio X10/1 genome and presented a high level of sequence diversity (**Figure 1a**). Sequence strand switches often delimited the surface molecule tandem arrays. Commonly, these arrays had two to four complete copies immediately followed by two or more truncated copies with motifs similar to the complete gene. The intergenic spaces between arrays were rich in simple and low complexity repeats with no identifiable regulatory elements. The VIPER and L1Tc retrotransposon elements, in clusters of two to four copies, were found in the proximity of, or inside, tandem arrays containing trans-sialidases, mucins and MASP genes. As the surface molecule genes are known to evolve rapidly and be highly variable^22^, the enrichment of VIPER and L1Tc elements in these regions supports the hypothesis that they may be involved in generating new surface molecule gene variants via recombination mediated by sequence homology.

Both, Ser/Thr kinases and DEAD-box RNA helicase genes were found at both extremes of 34 (10.81 %) trans-sialidase arrays located in pseudomolecules 1, 2 and 8. Searches against the RFAM database identified 1,618 small RNAs in the TcI Sylvio X10/1 genome. These were mostly ribosomal RNAs with the 5S rDNA subunit being the most common (31.9 %) followed by ACA Box snoRNAs (30.9 %), SSU rDNA (12.2 %) and LSU rDNA (10.2 %) subunits. We also found hits to telomerase RNA component (TERC), Catabolite Repression Control sequester (CrcZ), Protozoa Signal Recognition Particle RNAs, spliceosomal RNA subunits and miRNAs. The putative miRNAs identified in Sylvio X10/1 belong to the MIR2118 and MIR1023 families, previously not found in protozoan parasites. The functional relevance of these predicted small RNAs will need to be further validated *in vitro*. The miRNA segments were located in both strands within 1 Kb of genes coding for DEAD-box RNA helicases surrounding surface molecule gene tandem arrays.

### Genomic variation within the *Trypanosoma cruzi* TcI clade

Intra-TcI genomic diversity was examined among 34 samples from six countries: United States, Mexico, Panama, Colombia, Venezuela and Ecuador, derived from a range of triatomine vectors and human patients of different clinical stages (**Table 2** and **Supplementary Table 1**). Our hybrid variant calling strategy allowed us to identify genomic variants in the core and multigenic-families clusters in a reliable fashion (See methods).

**Table 2:** Genomic variants identified among the *Trypanosoma cruzi* TcI isolates.

| GROUP | SNPs | INDELs | DELETION | DUPLICATION | TRANSLOCATION |
| --- | --- | --- | --- | --- | --- |
| Colombia* | 158565 | 59520 | 439 | 1231 | 4140 |
| Colombiana** | 105023 | 30697 | 23 | 86 | 273 |
| Venezuela | 77232 | 70086 | 43 | 183 | 614 |
| Ecuador | 122122 | 84201 | 40 | 164 | 354 |
| Panama | 620499 | 238833 | 225 | 605 | 2060 |
| Texas | 101771 | 78499 | 69 | 303 | 978 |

\* - FcHc and CG clones from Colombia
\*\* - TcI Colombiana strain

A total of 1,031,785 SNPs and 279,772 INDELs shorter than 50 bp were called for all TcI sequenced isolates relative to the Sylvio X10/1 genome. INDELs presented an average density of 5.3 variants per Kb and SNPs 24.1 variants per Kb. An individual *T. cruzi* TcI isolate was found to contain an average of 61,000 SNPs and 6,820 INDELs with a density of 31.8 variants per Kb. However, these measures fluctuated depending on the geographical and biological source of the sample. Core regions had an average SNP density of 0.4 variants per Kb, in contrast with multigene-families clusters where values of 10 variants per Kb were found. It was not surprising that the bulk of the genomic variants were located in the multigenic families clusters regions in all the isolates, with fewer differences in the core regions. Although several studies using single gene markers have identified heterogeneity in the TcI clade^15,24^, the extent of this variation has not been previously assessed genome-wide. The majority of INDELs (96 %) were found in intergenic or noncoding regions, and 81 % of those were located in subtelomeric regions. INDELs within coding sequences were exclusively found to cause frameshifts turning the affected coding sequence into a pseudogene. This distribution of INDELs is a genomic signature that has been associated with non-allelic homologous recombination due to unequal crossing over^25^ or microhomology-mediated end joining^26,27^ (**Table 3**). Short insertions were more prevalent than short deletions, a pattern common to all the analysed TcI genomes when compared to Sylvio X10/1. In the subtelomeric regions, short insertions (1 - 3 bp) occurred within the upstream and downstream portions of the coding sequences and usually involved the addition of one or more cytosines or guanines. Deletions of 1 bp indicating the removal of an adenine or thymine were also observed within these regions, but at a lower frequency. Longer deletions (5 - 20 bp) and insertions (8 - 10 bp) were observed within trans-sialidases, Retrotransposon Hot Spot (RHS), pseudogenes and, at a lower frequency, L1Tc retroelements.

**Table 3:**
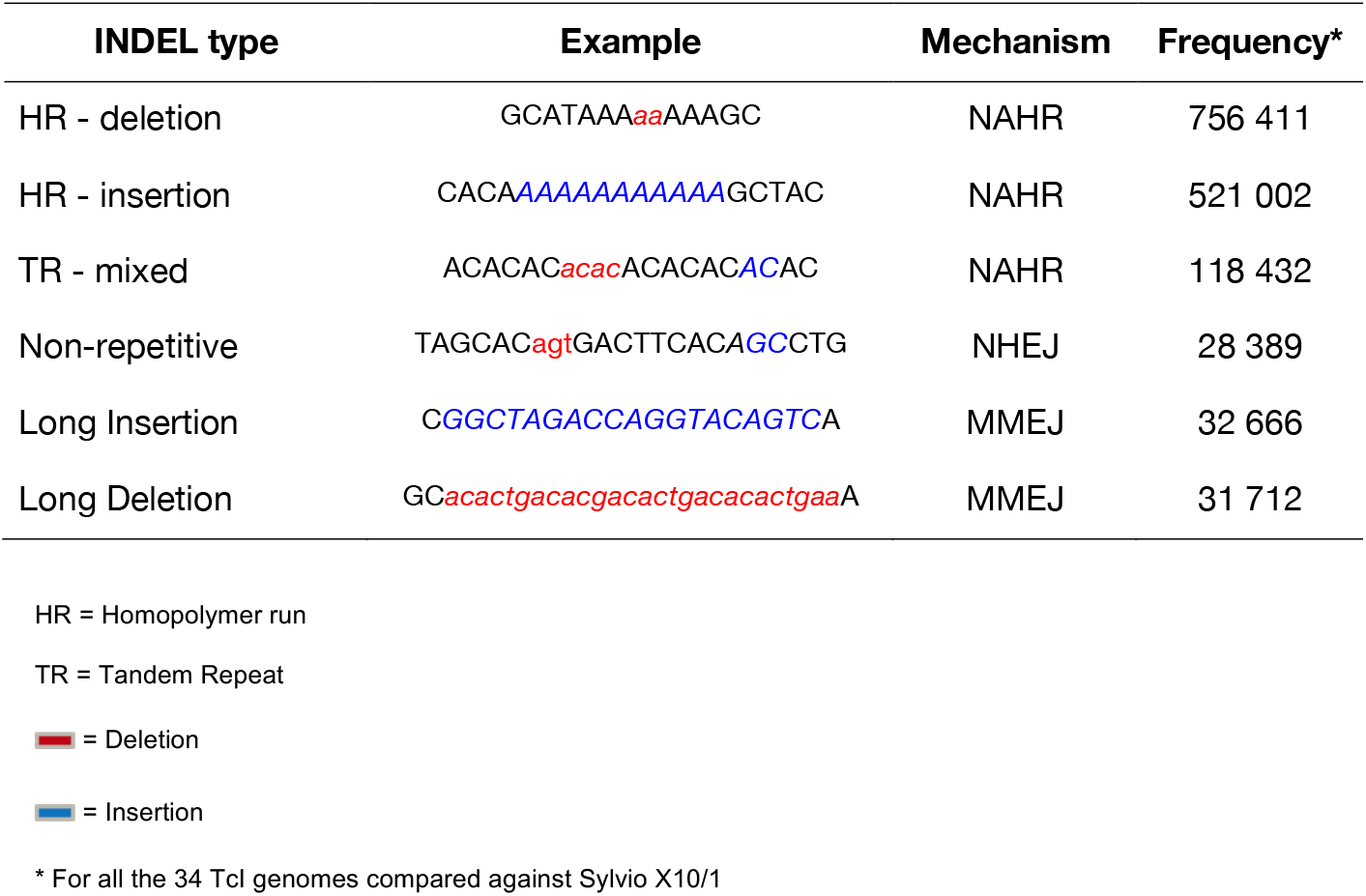
Patterns of INDELs and their associated mechanisms of origin.

### Population genomics of the *Trypanosoma cruzi* TcI clade

We used the short genomic variants to analyse the population genomics of the *T. cruzi* TcI clade, and where possible taking into account the different sample sources (insect vector or human host), clinical outcome of the infected patients and geographic locations (**Supplementary table 1**). This sampling strategy allowed comparison of parasite population structure in different environments. Interestingly, a Bayesian PCA analysis using INDELs and an IBD-based hierarchical clustering using only SNPs for all the samples showed a mostly geography-specific population structure (**Figure 3a** and **Supplementary Figure 3**).

**Figure 3:**
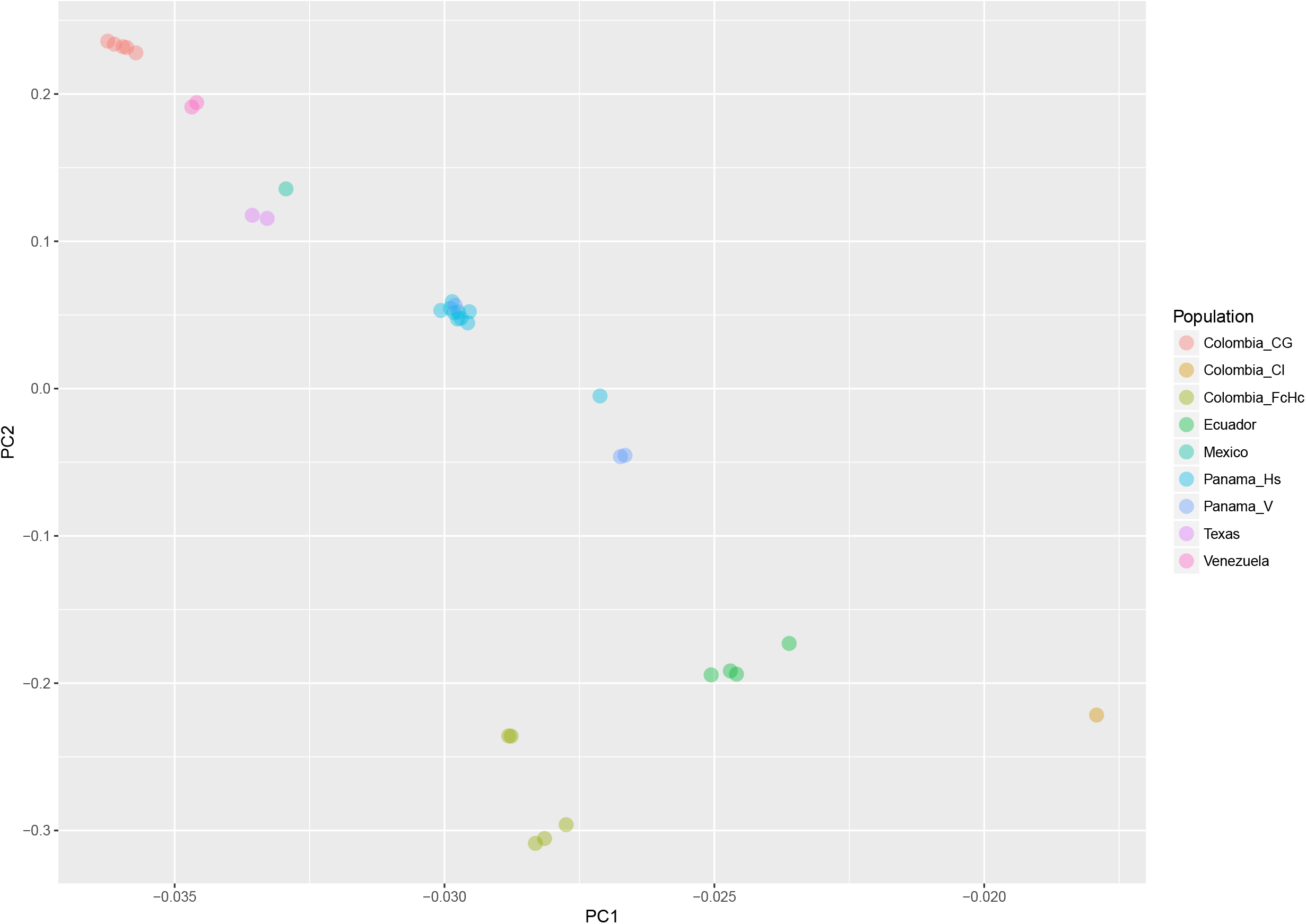
**a**) Linkage disequilibrium matrix (r2) of chromosome 2 for the Colombian CG and FcHc clones. LD values range from 0 (recombination) to 1 (no recombination). **b**) Genome-wide *Fst* distribution in 10 Kb bins displaying a state of panmixia for the CG clones and moderate genetic differentiation in the FcHc clones, yellow dots represent outlier bins. c) Distribution of subtelomeric *Tajima’s D* selection test in both groups displaying overall balancing selection (D > 0) in these regions for both clones.

The analysis of the variation between two Colombian TcI isolates made it possible to compare parasites from a HIV-positive patient with fatal cardiomyopathy (CG) and from an acute chagasic patient infected by oral transmission (FcHc). For each strain, replicate clones from the original sample were isolated and cultured under the same conditions, and five of the replicates from each sample were sequenced in a single Illumina HiSeq 2500 run and 158,565 well-supported SNPs were called. Using this set of SNPs we calculated global and per-site population genetic statistics. These samples displayed distinctive behaviour in a global analysis of genomic diversity by separating into two well-defined clusters, as can be seen in **Figure 3b**. Linkage Disequilibrium (LD) analyses were performed genome-wide for both groups using the r2 statistic; revealing a fluctuating pattern of LD across the entire genome with large blocks of low r2 values - implying a recombinatorial process - present at distinctive chromosomal locations that were specific to each group of clones. Particularly, CG clones had less genetic diversity than FcHc clones (**Figure 4a**) and displayed a trend towards LD, whereas FcHc clones presented more dynamic LD pattern. Values of r2 near zero were more common in LD sliding windows containing genes coding for surface molecules and r2 values closer to one were present exclusively in core regions rich in housekeeping genes, indicating that these regions are more stable. For the CG and FcHc clones we calculated a global Fixation index (*Fst*) value of −0.9377958 and −0.1162212 respectively (**Figure 4b**). These values are consistent with genetic differentiation in recombination hotspots in the subtelomeric regions. The global Tajima’s D value for the CG clones was 1.373 and 0.9906 for FcHc clones, suggesting the presence of multiple alleles at variable frequencies in both populations (**Figure 4c**). This pattern was more evident in the subtelomeric regions, which is consistent with balancing selection of surface molecules.

**Figure 4:**
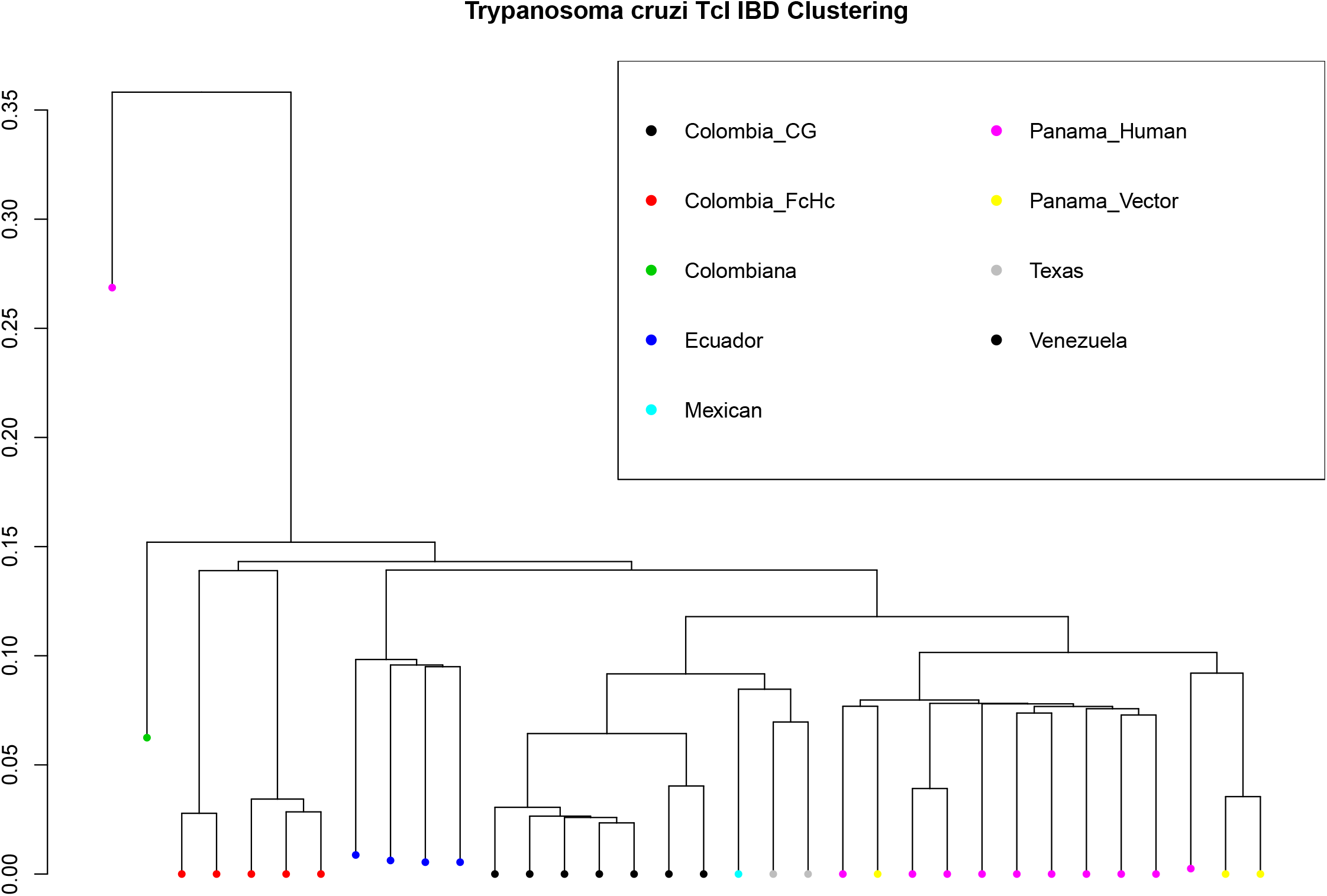
**a**) Proposed mechanism of inter-chromosomal recombination between gene tandem arrays for the generation of antigenic diversity. Here, VIPER retrotransposons (green) are surrounding a tandem array, shown in the direction of transcripton, containing trans-sialidases (blue) and pseudogenes (red) in pseudomolecules 1 and 8. The rearranged tandem array is shown below displaying the merger of genes and pseudogenes to form a longer coding sequence. **b**) Detail of simple repeats and retrotransposons within a trans-sialidase tandem array in pseudomolecule 1.

Analyses of genomic variation between samples isolated from humans and vectors from Mexico, Panama and Ecuador revealed that the global genetic differentiation among samples isolated from vectors was *Fst* = 0.1289547 whereas for samples isolated from humans the observed was *Fst* = −0.05521983. The patterns of linkage disequilibrium between human and vector derived isolates were similar to those observed in the Colombian clones. Estimates of the Tajima’s D statistic revealed a distinctive pattern of selection between the two groups. Balancing selection was detected specifically in regions containing tandem gene arrays coding for surface molecules in all the samples derived from vectors, regardless of their geographical origin; whereas selective sweeps were present in the same regions in human-derived samples. Large genomic areas (> 50 Kb) containing surface molecule genes displayed negative Tajima’s D values in human-derived isolates, in contrast with the pattern observed in vector-derived isolates with long genomic stretches (> 70 Kb) of positive Tajima’s D values and short genomic blocks (< 5 Kb) with negative values.

### Genome Structural Variation

Genomic structural variants, such as deletions, tandem and interspersed duplications, genomic inversions and chromosomal break-ends, were observed ubiquitously throughout the genomes of the analysed TcI strains. The most common type of intrachromosomal structural variant observed was tandem duplications followed by deletions larger than 50 Kb (**Table 2**). Chromosomal break-ends, similar to the unbalanced chromosomal translocations observed in higher eukaryotes, were the most abundant type of structural rearrangement and they were only present in genomic regions that were statistically enriched with retroelements and simple repeats. These areas presented a conserved pattern: they contained surface molecule gene tandem arrays and their breakpoints were composed of simple repeats and retrotransposons of the VIPER and L1Tc class.

These events were between 20 - 150 Kb in length and contained fragments or even complete coding sequences for surface molecule genes, such as trans-sialidases, mucins and MASP genes and surface glycoproteins (gp63/gp85). Housekeeping genes seemed to have not been affected by these genomic rearrangements. The breakpoints were composed of simple repeats, retrotransposons or both. Rearrangements affecting gene tandem arrays generated longer coding sequences by superimposing fragments - or the entire coding sequence - on genes of the same family located in a different genomic location. For instance, the Colombian isolates generated longer trans-sialidase genes by moving coding sequences from pseudomolecule 1 to pseudomolecule 8, while Texas isolates recombined trans-sialidases between pseudomolecule 16 and pseudomolecule 21. In this way, surface molecule genes were merged with another member of the same gene family - or a pseudogene - resulting in a new mosaic gene sequence (**Figure 5a**).

**Figure 5:**
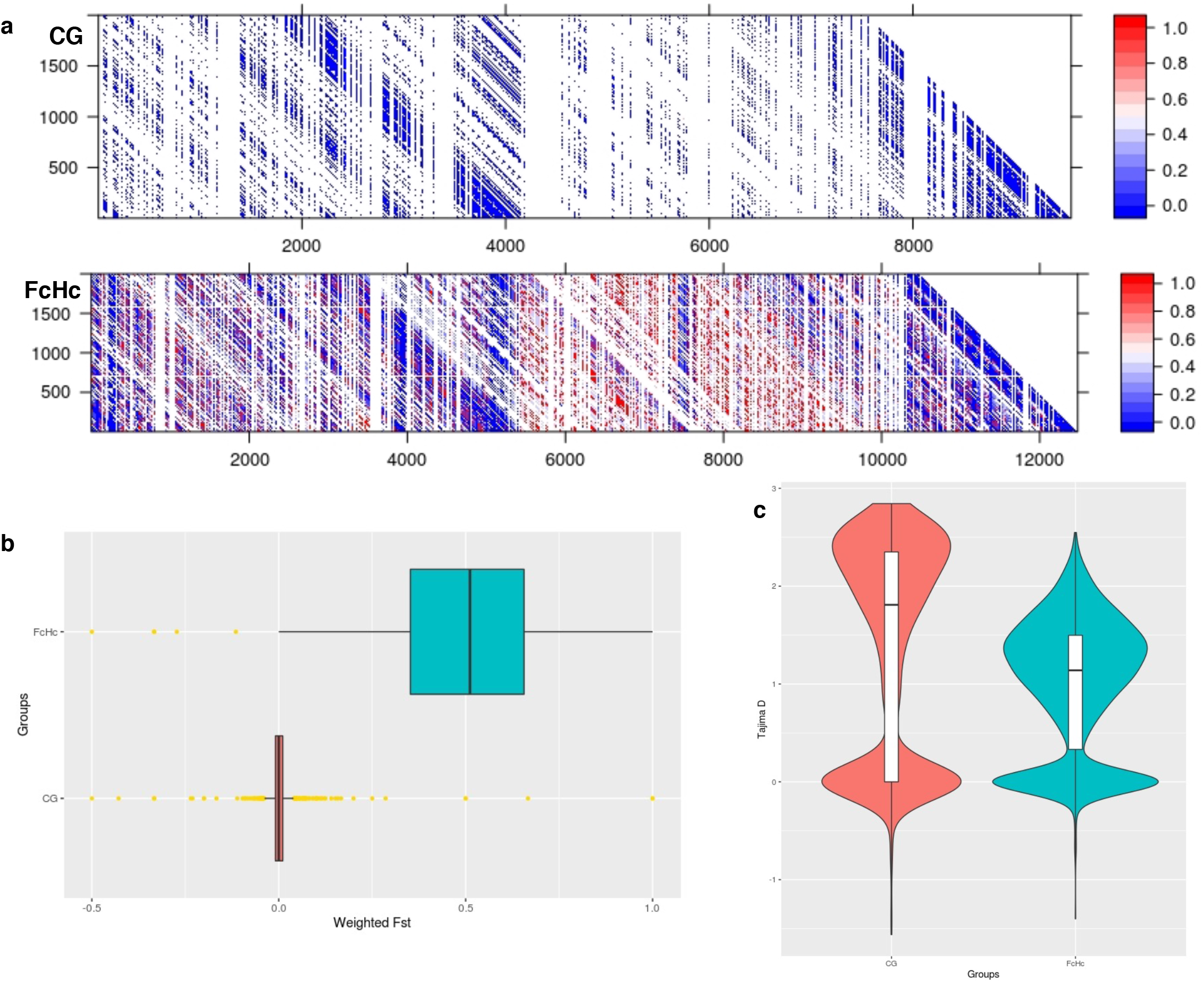
Distribution of CNV changes in chromosome 3 of the Colombian **a**) CG clones and **b**) FcHc clones. Black lines represent the reference genome sequence and cyan lines represent the sample under study. Each sliding window for CNV evaluation is represented as a dot. A drastic change in CNV can be noted in the FcHc clones whereas the CG clones seem to be less affected.

Retroelements could be found within or near genomic regions containing surface molecule gene tandem arrays (**Supplementary Tables 2, 3** and **4**) and L1Tc fragments or their entire sequence were also included in the rearranged region in all the observed translocation spots, where they were inserted into regions containing simple repeats composed by AT dimers (**Figure 5b**).

Multiple examples of the generation of new surface molecule gene variants were identified in TcI from diverse sources. It therefore appears that the parasite uses specific molecular mechanisms of recombination that can rapidly generate surface molecule diversity, allowing it to increase the genomic plasticity required to adapt to changing environments and evade immune responses during short and long-term infections in various host species.

The sizes of the tandem duplications ranged from 6 - 75 Kb and mainly involved tandem arrays coding for surface molecules, mostly trans-sialidases and mucins, but also Disperse Gene Family 1 (DGF-1) and several hypothetical proteins. The breakpoints of these duplications were surrounded by simple repeats and retroelements in subtelomeric regions. A tandem duplication event could involve between four and 25 copies of a specific gene when in the subtelomeric regions, whereas in core regions the number was between two and eight. We observed that large deletions occurring in subtelomeric regions were surrounded by simple repeats of the type (T)n and (AT)n and retrotransposons of the L1Tc class, containing surface molecule gene tandem arrays. Deletions in these genomic regions tended to be shorter (4 - 12 Kb) and sample specific.

### Distribution of Copy Number Variation (CNV) within the TcI clade

CNV varied extensively between *T. cruzi* TcI strains. Most notably, among the Colombian strains, isolates derived from the same sample presented different gene copy numbers. There have been previous attempts to assess CNV in the *T. cruzi* genome^28^, but these studies were performed using DNA tiling microarrays with probes designed using the TcVI CL Brener strain assembly, in which subtelomeric regions are essentially absent.

The distribution of CNV in the genomes of the studied TcI samples was isolate-specific, and involved segments of an average size of 5 Kb. In the samples analysed in our study we observed blocks of segmental CNV within a chromosome with a pattern that was unique to each sample. Notably, the Colombian clones presented individual profiles of CNV (**Figure 6a** and **6b**) despite being derived from the same clinical isolates.

Sequence blocks affected by segmental CNVs contained retrotransposons of the VIPER and L1Tc class, as well as surface molecule genes surrounded by simple repeats. The isolate-specific nature of these CNV events demonstrates the high level of within-clade diversity of the TcI samples. The distribution of CNV across the *T. cruzi* genome reinforces the dynamic nature of the multigene families clusters and the surface molecule gene families. As discussed below, it is important to note the association of structural and copy number variation with the presence of retrotransposons and simple repeats and their putative involvement in the generation of novel sequence variants via recombination.

## DISCUSSION

Complete reconstruction of the *T. cruzi* genome to encompass the subtelomeric regions and multigene families’ clusters, has proved to be difficult to achieve using short reads, due to sequencing library preparation biases and a genome architecture that is rich in long stretches of simple repeats, large repetitive gene families and multiple retrotransposons. Here we have used long PacBio sequencing reads to provide the most complete genome sequence of a *T. cruzi* strain to date. This has allowed us to perform the first detailed analyses of the repertoire of complex genes families that encode cell surface molecules, considered to be involved in cell invasion and evasion of the host immune response. We have shown the duality in the organisation of the parasite genome, comprised of a core genomic component with few repetitive elements and a slow evolutionary rate, resembling that of other protozoa, and a contrasting, highly plastic multigene families clusters encoding fast evolving surface antigens, with abundant interspersed retrotransposons. The structural changes that generate and maintain diversity in *T. cruzi* surface molecules have certain mechanistic parallels in other protozoa such as those recently described in *Plasmodium falciparum*^28^, but differing from the shorter, less repetitive genome of the non virulent, human-infective *Trypanosoma*^22^.

Early studies of the genetic diversity of *T. cruzi* using geographically disparate sampling and restricted comparisons of genetic diversity suggested a clonal population structure^29,30^; however, population genetics with an expanded set of markers have now challenged this view^15,31,32^. Nevertheless, there are still conflicting views as to which model best describes the population structure of *T. cruzi*^33,34^. The newer Sylvio X10/1 genome sequence will now enable extensive genome-wide comparative population genomics analyses, which may shed light on this issue. Comparative analyses of 34 *T. cruzi* isolates and clones from the TcI clade suggest many recombination events and population indices, normally associated with genetic exchange between strains, which are more likely to be caused by the extensive repeat-driven recombination in the subtelomeric regions. The extent of variation in the multigene-families clusters rich in surface antigen coding genes and the geographical clustering of strains based on geographic distribution indicates active, on-going adaptation to host and vectors. This need for phenotypic - and thus genomic - versatility may impel the active generation of sequence diversity in *T. cruzi*. Further analyses of the evolution of multigene families’ clusters will yield much more detailed understanding of diversity within and between the six currently recognised genetic lineages of *T. cruzi*^23^.

We have shown how the genome architecture and dynamic multigene families’ clusters of *T. cruzi* may provide a mechanism to rapidlygenerate sequence diversity, required to escape the host immune response and adapt in response to new environments. It is the striking richness in simple repeat, retrotransposons and motif conservation in the multigene families clusters that renders these genomic areas susceptible to structural change, similar to yeast and other pathogens (Weatherly 2016 - PMID: 27619017)^35,36,44^. Retrotransposons have been associated with the generation of complexity in genomic regions in mammals and plants and with control of gene expression^36,37^. In the case of *T. cruzi*, they appear to generate novel variants via mechanisms that exploit sequence homology. The presence of the simple repeats and retrotransposons near surface molecule coding genes provides the microhomology for both mechanisms to operate in such regions. Besides retrotransposons, the modular structure of the multigene families MASP and Trans-sialidase, where different genes share conserved motifs, could also provide microhomology needed for this homologous recombination (EL Sayed 2005 PMID: 16020725, Weatherly 2016 PMID: 27619017). Our analysis of INDELs and chromosomal breakpoints in the subtelomeric regions confirmed that a mechanism similar to NAHR or MMEJ operates as source of sequence diversity, for example by transposition of trans-sialidase genes or pseudogenes to produce new sequence mosaics. The required recombination machinery is conserved in *T. cruzi*^38^. Furthermore, these mechanisms would explain the high level of pseudogenisation observed in *T. cruzi*.

Retrotransposons were first reported from *T. cruzi* in 1991^39^. The presence of these elements may also partly account for the previously reported widespread observation of copy number variation in different *T. cruzi* strains^27^. Thus, we find that repeats near the surface molecule genes appear to drive recombination in *T. cruzi*. The apparent inability of *T. cruzi* to condense chromatin may facilitate transposition in a stochastic fashion, facilitating generation of sequence diversity in exposed regions of the genome. A similar process has been described in the neurons of higher eukaryotes^40^ but not in any other unicellular organism. Retrotransposons may also have an important role as gene transcription regulators: they may either silence or promote gene expression, due to their susceptibility to DNA methylation or by providing potential binding sites respectively, as it has been observed in previous works^41^. This lack of a well-defined transcriptional regulation machinery in the *T. cruzi* genome may suggests a link to the requirement for retrotransposon closely associated with gene tandem arrays.

## CONCLUSION

Here we have sequenced and assembled the complete genome of a *Trypanosoma cruzi* TcI strain. This has enabled the first resolution of the complex multiple gene families that encode *T. cruzi* surface molecules, and provided a basis for *T. cruzi* population genomics. We discover an extraordinary concentration of retrotransposons among the multigene families’ clusters and indications of repeat-driven recombination and generation of antigenic diversity, providing the mechanisms for *T. cruzi* to evade the host immune response, and to facilitate the adaption to new host and vectors. This genome will provide an invaluable resource to facilitate the prospective discovery of novel drug targets and vaccine candidates for Chagas disease.

## METHODS

### Genome Sequencing and Assembly

*Trypanosoma cruzi* Sylvio X10/1 strain was isolated from an acute human case of Chagas disease in Brazil^11^. Total genomic DNA of this TcI strain was obtained from culture epimastigotes as formerly described^11^ and used to produce PacBio CCS data according to standard protocols from the Genomic Facility of Science for Life Laboratory (Sweden) and Pacific Biosciences (USA). Genomic DNA was sequenced to a depth of 210X using the PacBio platform, supplying raw reads with an average length of 5.8 Kb. These reads were corrected by means of the PBcR v8.3 pipeline with the MHAP algorithm^42^ using the auto-correction parameters described to merge haplotypes and skipping the assembly step, producing a total of 1,216 contigs (NG50 = 62 Kb). Later, the assembly was scaffolded using the corrected PacBio reads with the SSPACE-Long scaffolder yielding 310 scaffolds (NG50 = 788 Kb); 118 gaps were filled using Illumina reads with GapFiller and corrected PacBio reads with PBJelly2. Finally, the core regions of these scaffolds were aligned against the core regions of the TcVI CL Brener reference genome using ABACAS (http://abacas.sourceforge.net), producing 47 pseudomolecules. The quality of the new assembly was assessed with FRC_bam with the Illumina paired end reads generated at the Genomic Facility of Science for Life Laboratory (Sweden) using the same genomic DNA extraction used for PacBio sequencing.

### Annotation of the *Trypanosoma cruzi* Sylvio X10/1 Genome

The genome sequence was annotated using a new kinetoplastid genome annotation pipeline combining homology-based gene model transfer with *de novo* gene prediction. To allow for the sensitive identification of partial genes, input sequences were split at stretches of undefined bases, effectively creating a set of ‘pseudocontigs’, each of which does not contain any gaps. Gene finding was then performed on both the original sequences and the pseudocontigs using AUGUSTUS, which also calls partial genes at the boundaries of each pseudocontig. AUGUSTUS models were trained on 800 genes randomly sampled from the 41 Esmeraldo-type (TcII) *T. cruzi* CL Brener chromosomes in GeneDB. Protein-DNA alignments of reference proteins against the new *T. cruzi* sequences, generated using Exonerate, were additionally used to improve the accuracy of the gene prediction. In addition, the RATT software was used to transfer highly conserved gene models from the *T. cruzi* CL Brener annotation to the target. A non-redundant set of gene models was obtained by merging the results of both RATT and AUGUSTUS and, for each maximal overlapping set of gene models, selecting the non-overlapping subset that maximizes the total length of the interval covered by the models, weighted by varying levels of *a priori* assigned confidence. Spurious low-confidence protein coding genes with a reading direction in disagreement with the directions of the polycistronic transcriptional units were removed automatically. The result of this integration process was then merged with ncRNA annotations produced by specific tools such as ARAGORN and Infernal. Finally, protein-DNA alignments with frame shifts produced by BLAST were used in a computational approach to identify potential pseudogenes in the remaining sequence.

Downstream of the structural annotation phase, gene models were automatically assigned IDs and further extended with product descriptions and GO terms, both transferred from CL Brener orthologs and inferred from Pfam protein domain hits and represented as feature attributes or Sequence Ontology-typed subfeatures tagged with appropriate evidence codes. This annotation pipeline has been implemented in the Companion web server. The assembled genome was scanned for small RNAs using INFERNAL against the curated RFAM database using cmsearch with a minimum e-value of 1e-10, a GC-bias of 0 and a minimum alignment length of 10 nt. This annotation process has been implemented into the web-based annotation pipeline COMPANION^43^ from the Wellcome Trust Sanger Institute.

Repetitive sequences were annotated using RepeatMasker with the NCBI+ search engine and LTRHarvest. Using the genomic coordinates of the repetitive elements, the genome was split in windows of 10 Kb to identify VIPER and L1Tc retroelements adjacent to surface molecule genes (i.e: trans-sialidases, mucins and MASP). A one-sided Fisher’s exact test was used to evaluate if the retroelements were enriched in genomic segments containing surface molecule genes.

### Identification of Single Nucleotide Polymorphism (SNP) and Insertion/Deletion events (Indels)

An improved short-read mapping strategy was used to assign the reads to their target sequences with high accuracy, especially in regions rich in simple and low complexity repeats, by taking advantage of the statistical read placement implemented in the Stampy read mapper to accurately call genomic variants from the mapped reads. Reads from all 34 *T. cruzi* TcI isolates (SRA BioProject accession number: PRJNA325924) were mapped against the assembled *T. cruzi* Sylvio X10/1 genome using a two-step mapping process to improve the mapping of Illumina data to highly repetitive regions: First, reads were mapped using BWA MEM with default parameters; later, the BAM file produced by BWA was remapped with Stampy (v1.23) using the *--bamkeepgoodreads* option. The final mapping file was sorted and filtered for PCR duplicates using Picard Tools v1.137. Variants were called using FreeBayes with a minimum per-base quality of 30, minimum mapping quality of 30 and minimum coverage of 15 bases. Variants that were found in a potentially misassembled region were excluded from the analysis. Additionally, genomic variants were called using FermiKit - which is an assembly-based variant caller - to validate the genomic variants observed in subtelomeric regions. A consensus of the two methods was used as a final set of variants for downstream analyses. Haplotypes were phased using Beagle r1399. The phased markers were used for downstream analyses with SNPrelate and VCFtools and the functional effect of the identified variants was predicted using SnpEff.

### Identification of Genomic Structural Variants (SV)

Genomic structural variants were identified whiting TcI isolates, using a consensus of different methods: Delly2, Lumpy, FermiKit and FindTranslocations (https://github.com/vezzi/FindTranslocations.git) using both raw reads and realigned BAM files. For each method, a SV muts had a depth of coverage > 10 reads and a mapping quality of > 30. Later, a consensus was created with all the SV that were supported by all the methods. SVs that were supported by FermiKit and at least one of the mapping-based methods were also included but labelled as ‘Low Confidence’. SVs identified by only one method were not included. Breakpoint analysis was done with custom Python scripts and their functional effect was predicted using SNPeff. Analyses of copy number variation (CNV) were done using the BAM files for each sample with the *Control-FREEC* package.

## Accessions

Whole genome sequence Genbank accession: ADWP00000000 (CP015651-CP015697). Reads for the 34 TcI isolates SRA accession: SRP076682.

## Authors contributions

CT-L and BA conceived and designed the study. CT-L designed and executed computational analyses. MY prepared Sylvio X10/1 genomic DNA for PacBio sequencing and performed manual annotation of surface molecule genes. JEC, AS, JDR, FG, SO-M, JAC, SMRT, HC, RG, KJ, MEB, PJH, KOM, MJG, BB provided genomic DNA for TcI isolates. JLR-C and DCB created chromosome maps for surface molecules. MAM, LAM, ML, JLR-C and ECG contributed to the interpretation of the results. CT-L, ECG, MAM, BA wrote the manuscript. All authors read and approved the final version of the manuscript.

## Acknowledgements

The authors wish to thank to John Kuijpers and Lawrence Hon from Pacific Biosciences (PacBio) for providing the data for the Sylvio X10/1 genome and preliminary assembly results. We would also like to acknowledge Eric Dumonteil, University of the Yucatan, Mexico, and Ed Wozniak, Texas Department of State Health Services, for past work on parasite collection. This research was funded by grants from the Knut and Alice Wallenberg Foundation, The Swedish Research Council and the European FP7 program. LAM was funded by a grant from the NIH; DCB, ECG and SMRT were funded by grants from CNPq (Brazilian Government Agency).

## SUPPLEMENTARY DATA

**Supplementary Table 1:** *Trypanosoma cruzi* strains used in this study.

| Strain | Country | Source | Clinical form | Sequencing coverage* |
| --- | --- | --- | --- | --- |
| H1a | Panama | Human | Chronic | 31 X |
| H2 | Panama | Human | Asymptomatic | 28 X |
| H3 | Panama | Human | Asymptomatic | 32 X |
| H4 | Panama | Human | No Disease | 29 X |
| H5 | Panama | Human | Chronic | 31 X |
| H6 | Panama | Human | Asymptomatic | 31 X |
| H7 | Panama | Human | Asymptomatic | 28 X |
| H8 | Panama | Human | No Disease | 32 X |
| H12 | Panama | Human | No Disease | 31 X |
| H14 | Panama | Human | Chronic | 32 X |
| H15 | Panama | Human | Chronic | 31 X |
| V1 | Panama | *P. geniculatus* | ND | 28 X |
| V2 | Panama | *R. pallescens* | ND | 25 X |
| V3 | Panama | *T. dimidiata* | ND | 28 X |
| TBM3324 | Ecuador | *R. ecuadoriensis* | ND | 28 X |
| TBM3406B1 | Ecuador | *R. ecuadoriensis* | ND | 25 X |
| TBM3479B1 | Ecuador | *R. ecuadoriensis* | ND | 29 X |
| TBM3519W1 | Ecuador | *R. ecuadoriensis* | ND | 28 X |
| X10462-P1C9 | Venezuela | Human | ND | 31 X |
| X12422-P1C3 | Venezuela | Human | ND | 29 X |
| Colombiana | Colombia | Human | ND | 30 X |
| CGI10 | Colombia | Human | Acute Co-infection | 30 X |
| CGI11 | Colombia | Human | Acute Co-infection | 30 X |
| CGI12 | Colombia | Human | Acute Co-infection | 30 X |
| CGI13 | Colombia | Human | Acute Co- infection | 30 X |
| CGI14 | Colombia | Human | Acute Co-infection | 30 X |
| CGI15 | Colombia | Human | Acute Co-infection | 30 X |
| FcHcl1 | Colombia | Human | Acute Oral | 30 X |
| FcHcl2 | Colombia | Human | Acute Oral | 30 X |
| FcHcl3 | Colombia | Human | Acute Oral | 30 X |
| FcHcl4 | Colombia | Human | Acute Oral | 30 X |
| FcHcl5 | Colombia | Human | Acute Oral | 30 X |
| H1b | Mexico | Human | Acute | 32 X |
| TD23 | Texas | *T. dimidiata* | ND | 30 X |
| TD25 | Texas | *T. dimidiata* | ND | 30 X |

NA = Not determined.
\* Illumina data

**Supplementary Figure 1:**
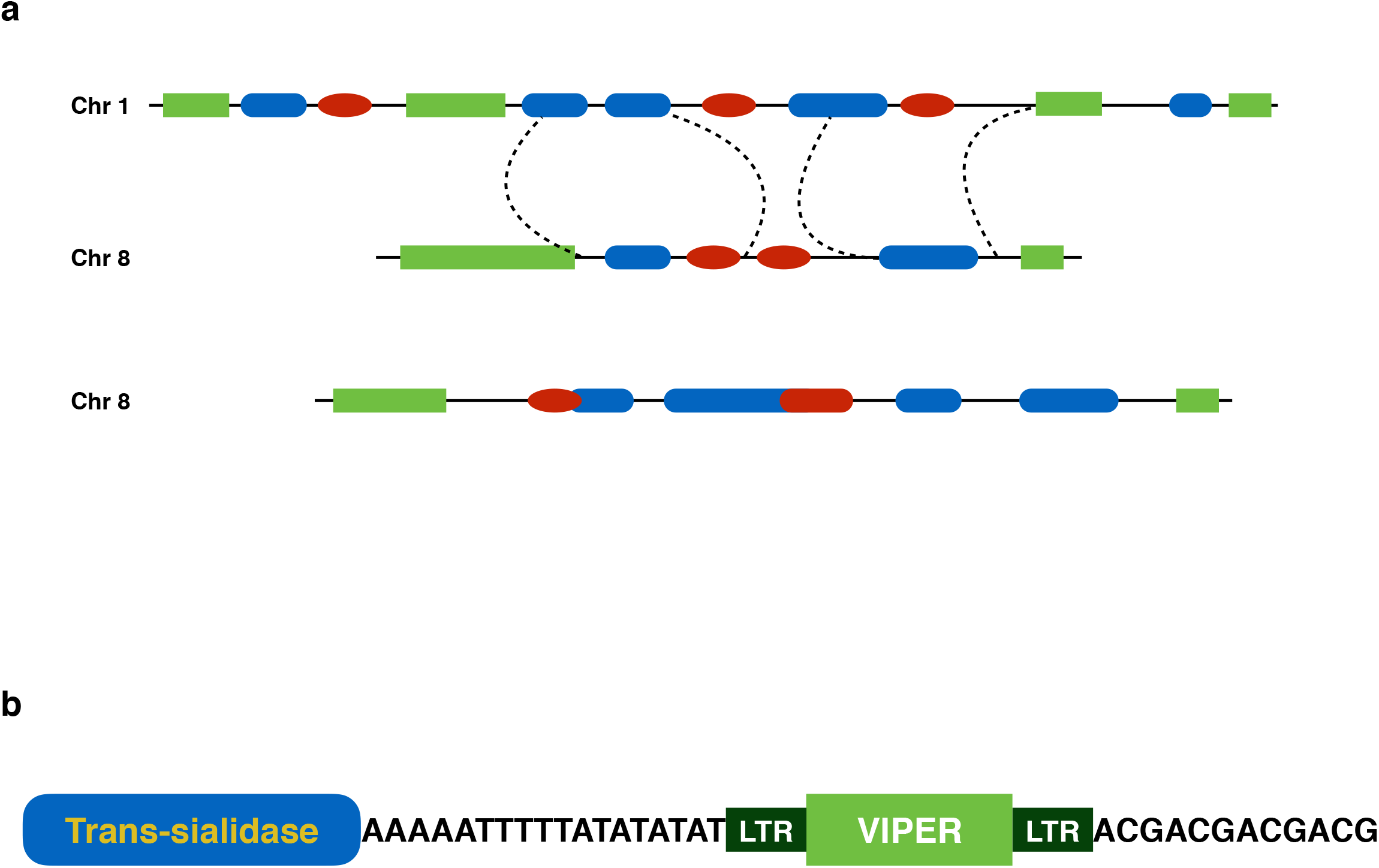
Distribution of coverage for **a**) PacBio and **b**) Illumina reads in a subtelomeric end on chromosome 8 showing a well supported genome assembly of these regions.

**Supplementary Figure 2:**
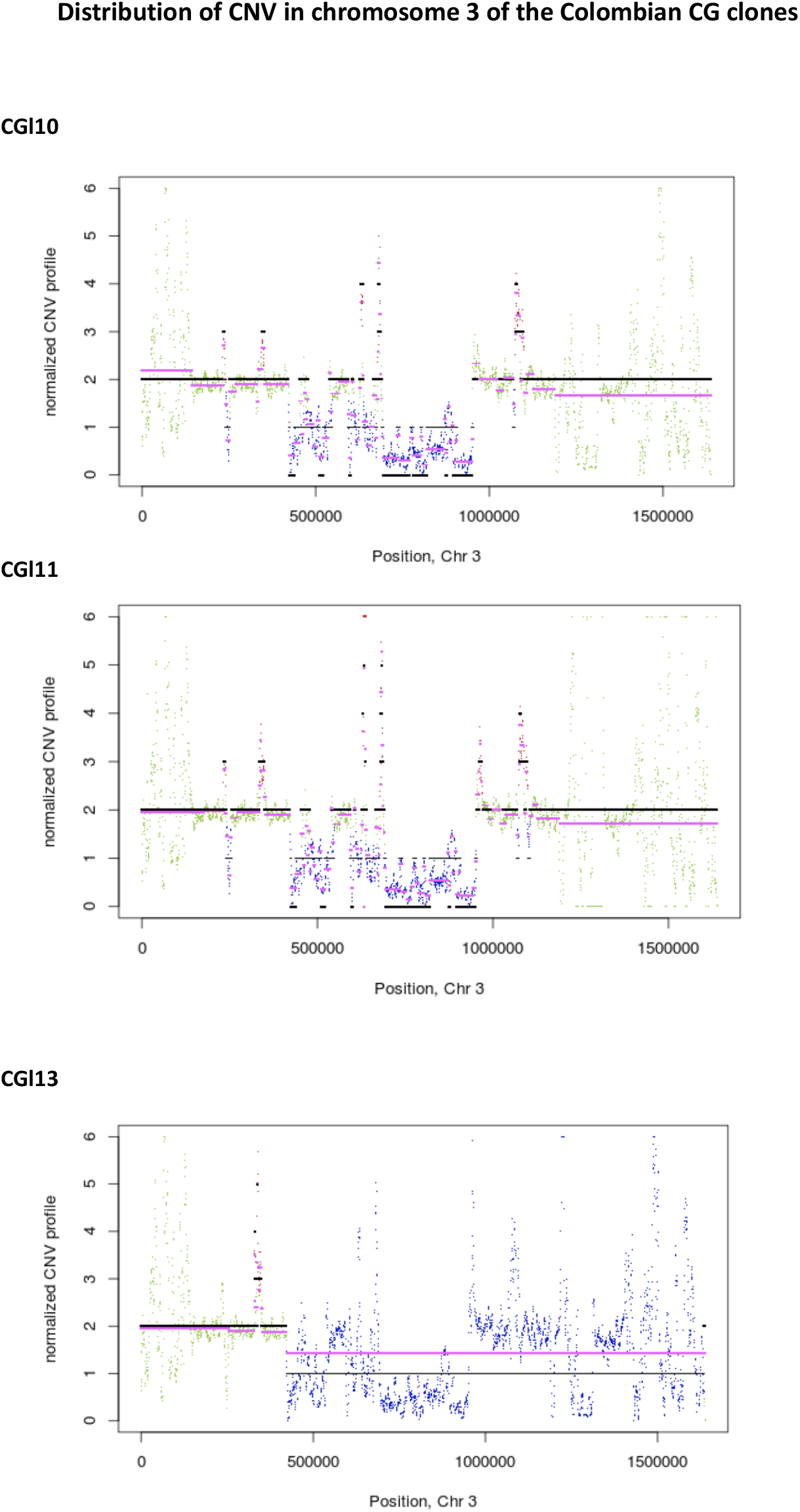
Genome-wide Neighbour Joining tree for trans-sialidases genes from the Sylvio X10/1 genome showing the high level of sequence diversity for these gene family.

**Supplementary Figure 3:**
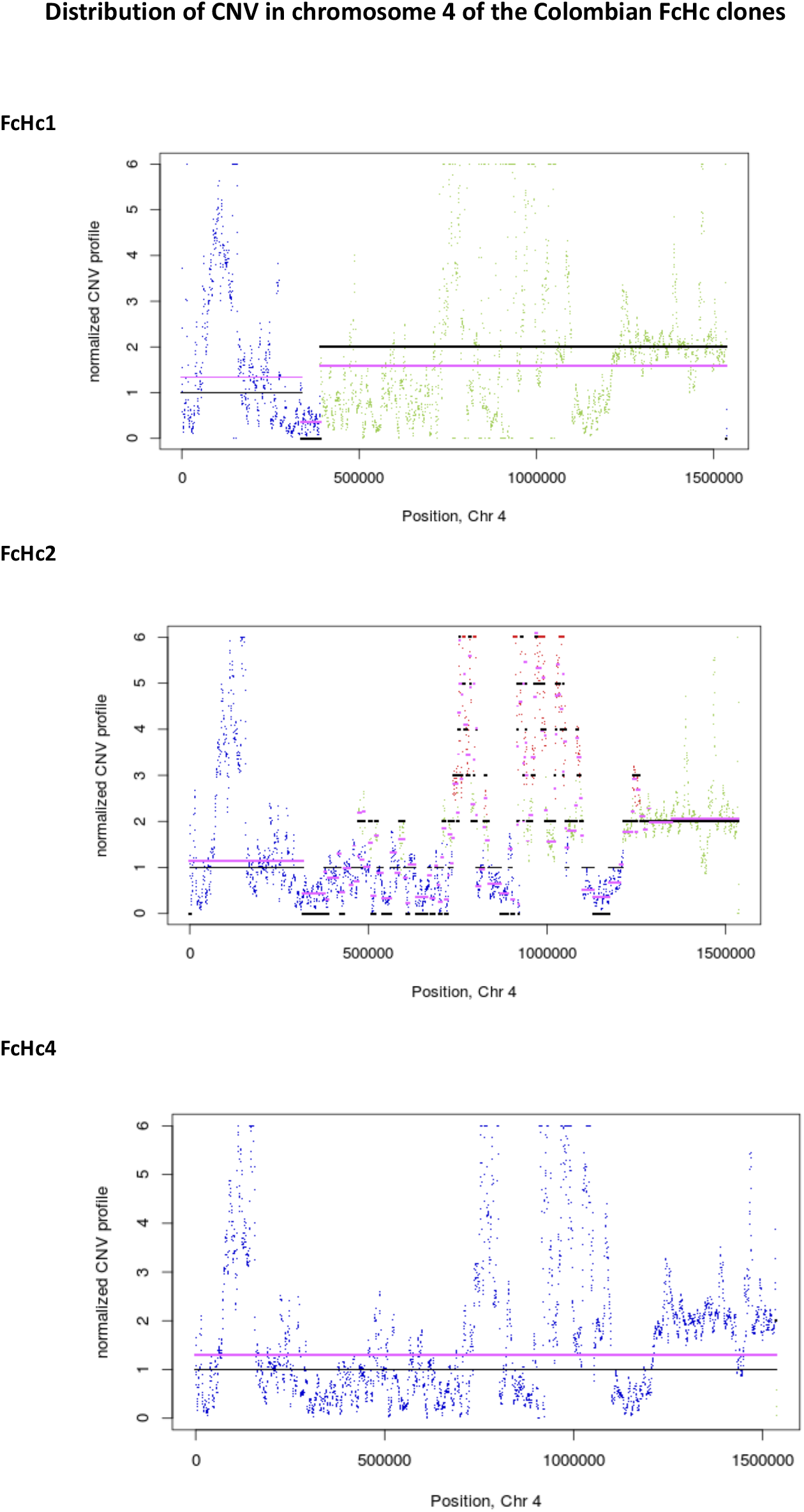
Principal component variability for the genotype diversity of the 34 TcI isolates.

